# mebipred: identifying metal-binding potential in protein sequence

**DOI:** 10.1101/2021.08.12.456141

**Authors:** AA Aptekmann, J Buongiorno, D Giovannelli, M Glamoclija, DU Ferreiro, Y Bromberg

## Abstract

Metal-binding proteins have a central role in maintaining life processes. Nearly one-third of known protein structures contain metal ions that are used for a variety of needs, such as catalysis, DNA/RNA binding, protein structure stability, etc. Identifying metal-binding proteins is thus crucial for understanding the mechanisms of cellular activity. However, experimental annotation of protein metal-binding potential is severely lacking, while computational techniques are often imprecise and of limited applicability.

We developed a novel machine learning-based method, *mebipred*, for identifying metal-binding proteins from sequence-derived features. This method is nearly 90% accurate in recognizing proteins that bind metal ions and ion containing ligands. Moreover, the identity of ten ubiquitously present metal ions and ion-containing ligands can be annotated. *mebipred* is reference-free, *i.e.* no sequence alignments are involved, and outperforms other prediction methods, both in speed and accuracy. *mebipred* can also identify protein metal-binding capabilities from short sequence stretches and, thus, may be useful for the annotation of metagenomic samples metal requirements inferred from translated sequencing reads. We performed an analysis of microbiome data and found that ocean, hot spring sediments and soil microbiomes use a more diverse set of metals than human host-related ones. For human-hosted microbiomes, physiological conditions explain the observed metal preferences. Similarly, subtle changes in ocean sample ion concentration affect the abundance of relevant metal-binding proteins. These results are highlight *mebipred’*s utility in analyzing microbiome metal requirements.

*mebipred* is available as a web server at services.bromberglab.org/mebipred and as a standalone package at https://pypi.org/project/mymetal/

## Introduction

Proteins bind a diverse set of metal ion-containing cofactors and sustain the functional requirements of life. Metal ions, *e.g.* iron, magnesium, copper, *etc*., and metal-containing ligands, *e.g.* heme and iron-sulfur clusters, participate in protein folding/stability [1], DNA replication [2–4], catalysis [5], redox chemistry [5] and many other cellular activities. Proteins could thus be described as sophisticated electron transfer nanomachines that depend on transition metal ions to perform their functions [6–8]. Of the proteins whose three-dimensional structure is available in the Protein DataBank (PDB) [9], roughly a third (49,996 of 152,346) are metal-binding proteins, a finding which may be somehow related to their high abundance in nature. However, only a small fraction of metal-binding protein sequences has been identified overall. The Swiss-Prot [10] database, for example, contains over half a million (564,638) manually-curated protein sequences, of which ~14% (94,720) are annotated as metal-binding; the binding activity of only a few of these (<1%, 4,251 proteins) has thus far been experimentally verified (Feb 2020). Furthermore, of the nearly 180 million proteins in TrEMBL, which are generated via translation of sequenced genome open reading frames and have no experimental annotations, only about five million sequences, i.e. less than 3%, are annotated as metal-binding [11].

Different levels of sequence redundancy in distinct databases may be an underlying cause for this discrepancy. However, another major reason is that we are still unable to accurately identify metal-binding proteins directly from their sequences and, in some cases, even from their high resolution structures [12]. Experiments, *e.g.* mass spectrometry [13] and crystallography [14], can detect protein-metal interactions, but these analyses are expensive and time-consuming, as well as error-prone for both technical and biological reasons. For example, cambialistic proteins can use metal cofactors interchangeably [15] and thus are likely to be misclassified when experimentally assessed for binding of specific metals. Similarly, some experiments include the use of non-native metals for technical and/or crystallization purposes [16], the loss of metal ion-binding ability/specificity in the process of protein purification [17], or even simply incorrect identification of the bound metals due to low experimental resolution [18]. Thus, metal-binding annotation for most protein sequences is lacking and, likely, only a small portion of extant metal-binding protein sequences has been identified.

There is no simple way to establish from sequence whether a protein binds a metal or not, but there have been multiple attempts to predict binding of single ion ligands (including metals) from protein structures. While a complete account of all relevant methods present in the literature is beyond the scope of this work (for a review see Lavechia et al [19]), here we highlight some trends in tool development.

Metal-binding sites in proteins frequently comprise a shell of hydrophilic residues that can be identified in the protein 3D structure [20]. For example, one algorithm [21] detects Ca^2+^ binding via identification of Ca^2+^ ion coordination by a layer of oxygen atoms supported by an outer shell of carbon atoms. Available structure-based methods use knowledge of hydrophilic shell residues to make predictions [20–24]. The main disadvantage of these approaches is that many such hydrophilic shells do not bind metals [25]. Additionally, structure-based methods are limited by the relatively small number of available protein structures (157,668; PDB, Feb 2020) [9]. However, when a protein structure is available, these methods often attain better performance than ones based on sequence alone.

To circumvent the limitation in the number of 3D structures, methods using homology modeling of proteins were developed. Early attempts at this type of prediction (*e.g.* MetSite [26]) had poor performance (58% precision at 28% recall). Overall, methods based on homology modeling tend to perform poorly when predicting sequences modeled with structural templates of less than 40% sequence identity; *e.g.* 42% precision at 65% recall [27]. Moreover, these methods attain a better performance when focusing on a single metal ion than when trying to describe binding of multiple ions, *e.g.* Liu et al calcium-binding site predictor (99% precision at 75% recall) [28] and Zhao et al zinc-binding predictor (90% precision at 72% recall) [29].

The computational prediction of metal-binding can be similar in essence to the prediction of other functional characteristics of proteins from sequence, *e.g.* mutation effects [30–32], residue importance [33], or subcellular localization [34]. Here, evolutionary profiles, predicted structure, physicochemical properties, and sequence descriptors are combined as features for machine learning. One such approach to the prediction of metal-binding [35] has attained fairly high accuracy (70% overall accuracy). Other methods combine structural and sequence features in a process known as “fragment transformation” [23, 35, 36]; e.g. Lin et al [23] report accuracies above 92%. Combining sequence, structure, and residue contact features in a random forest framework, the tool MetalExplorer [37] predicts the binding of eight metal ions. Performance across ions is varied, with a precision of 60% for recalls ranging from 59% to 88%.

There are also structure-independent (purely sequence-based) methods to predict metal binding. Function transfer by homology, *i.e.* the assumption that similar sequences perform similar functions, is one of the simplest ways to infer metal binding for protein sequences. Similarity is often established by alignment methods. However, it remains unclear that there is a well-defined alignment score cutoff for identifying functionally similar proteins [38]. Moreover, sequence similarity, or even well-characterized homology, may be misleading as homologs may evolve to bind different metals due to changing environmental pressures [39]. It is also possible to predict metal binding using sequence conservation of residues near those directly interacting with Zn^2+^, Cu^2+^, Fe^2+^, Fe^3+^, and Co^2+^ ions with a high accuracy [40]; proteins binding other ions were not identified using this method. Pattern recognition (*e.g.* Hidden Markov Models, HMMs, [41] and regular expressions, *e.g.* [42]) can also be used to expand the suspected set of metal binding sequences on the basis of remote homology. Unfortunately HMMs, designed to identify evolutionary conserved sequence patterns, are too specific and, thus, not well-suited for *de novo* metal binding prediction.

More complicated sequence-based metal-binding predictors often use machine learning techniques (e.g. neural networks [43] support vector machines (SVM) [44, 45], and random forests [46]). The performance these methods varies; *e.g.* Lin et al [47] reported high precision for all ions, albeit at recall as low as 35%. Combining different methods to identify specific residues involved in metal binding, *e.g*. Zn-binding cysteines and histidines, produced high accuracy [45, 48–50]. Note that while all of the above methods report good performance, we were unable to validate these reports using our own data as the webserver/standalone versions (where applicable) were nonfunctional and downloadable scripts absent.

Here we present *mebipred* (<u>metal</u>-<u>bi</u>nding <u>pred</u>ictor), a computational method for the prediction of protein metal binding potential based on sequence information alone. Our method is widely applicable because it doesn’t depend on the existence of a high resolution structure, has a better performance (average precision/recall of 95/78% at default cutoff) and is faster (17,000 sequences/minute) than existing sequence-based tools, and can be used to predict metal binding using whole protein sequences as well as short peptide fragments. The latter ability makes it potentially suitable for annotation of shotgun-sequenced unassembled metagenomic data/reads. *mebipred* is also alignment-free and thus useful for the analysis of newly identified proteins (with no known homologs). Finally, as mentioned previously, *mebipred* is the only currently publicly available method for sequence-based prediction of metal binding.

## Methods

### Datasets

We explored proteins binding Na, K, Ca, Mg, Mn, Fe, Cu, Ni, and Zn metal-containing ligands, regardless of their oxidation state (*e.g.* Fe2+ and Fe3+ are both in the Fe class) or context (*e.g.* Fe-containing hemes are in the same class as Fe ions). We retrieved all protein structures with these metal-containing ligands from the PDB (July 2019) and parsed them using the BioPython PDB module [51] (Supplementary Table 1). One naive approach to identify a set of metal-binding proteins is to compile all structures that have a metal ion. However, in the case of heteromers, *i.e.* protein complexes that contain multiple nonidentical chains, it is possible that only one of the chains binds the metal. We thus considered as metal-binding only the amino acid sequences/chains with at least one heavy atom within 5Å of the metal ion (METAL set). All other chains were included in the NO_METAL set, along with all PDB structures that contained no metals at all. Note that this criterion for the differentiation of metal-binding/nonbinding chains could lead to disagreement with existing metal-binding annotations.

### Feature extraction

To describe the proteins in our METAL and NO_METAL sets, we used only sequence-based features: 1) amino acid composition, 2) amino acid physicochemical properties, and 3) a count of the metal-binding amino acid 5mers.

1. *Amino acid composition*: for each of the 20 standard amino acids, the percentage of the type of amino acid in the entire sequence. *Total: 20 features.*
2. *Physicochemical properties of amino acids*: We used a set of nine amino acid properties as described in Li et al. [52]: hydrophobicity, hydrophilicity, number of hydrogen bonds, volume, polarity, polarizability, solvent accessible surface area, net charge index of side chains, and average amino acid mass. For each of the properties (Fig. 1), we clustered sequence residues into three categories, Low (L), Medium (M), and High (H), using “Jenks natural breaks” criterion [53]; this measure seeks to minimize each class’s average deviation from the class mean while maximizing each class’s deviation from the means of the other classes. For each category, we further calculated its *composition* (C), *i.e.* the fraction of amino acids in the sequence belonging to the category; *transitions* (T), *i.e.* for every pair of sequential amino acids, the number of transitions from one category to another divided by sequence length; and *distribution* (D), *i.e.* the sequence position of the amino acid corresponding to category’s first occurrence, 25% of occurrences, 50%, 75%, and last occurrence, divided by sequence length. *Total: 219 features.*
3. For each protein structure in the PDB, individually for every metal, we identified all five amino acid-long subsequences (5mers) within 5Å of the bound metal ion (Fig. 2; i.e. for heme this means within 5Å of the Fe atom) and counted the frequency of their occurrence in all complete PDB sequences. Every query sequence is then decomposed into 5mers (via sliding window of 1) and the 5mer feature is computed as the sum of all occurrences of the individual metal-specific 5mers in the PDB dictionary. We also included as a feature the sum of the frequencies across all ions. *Total: 11 features.*

**Figure 1:**
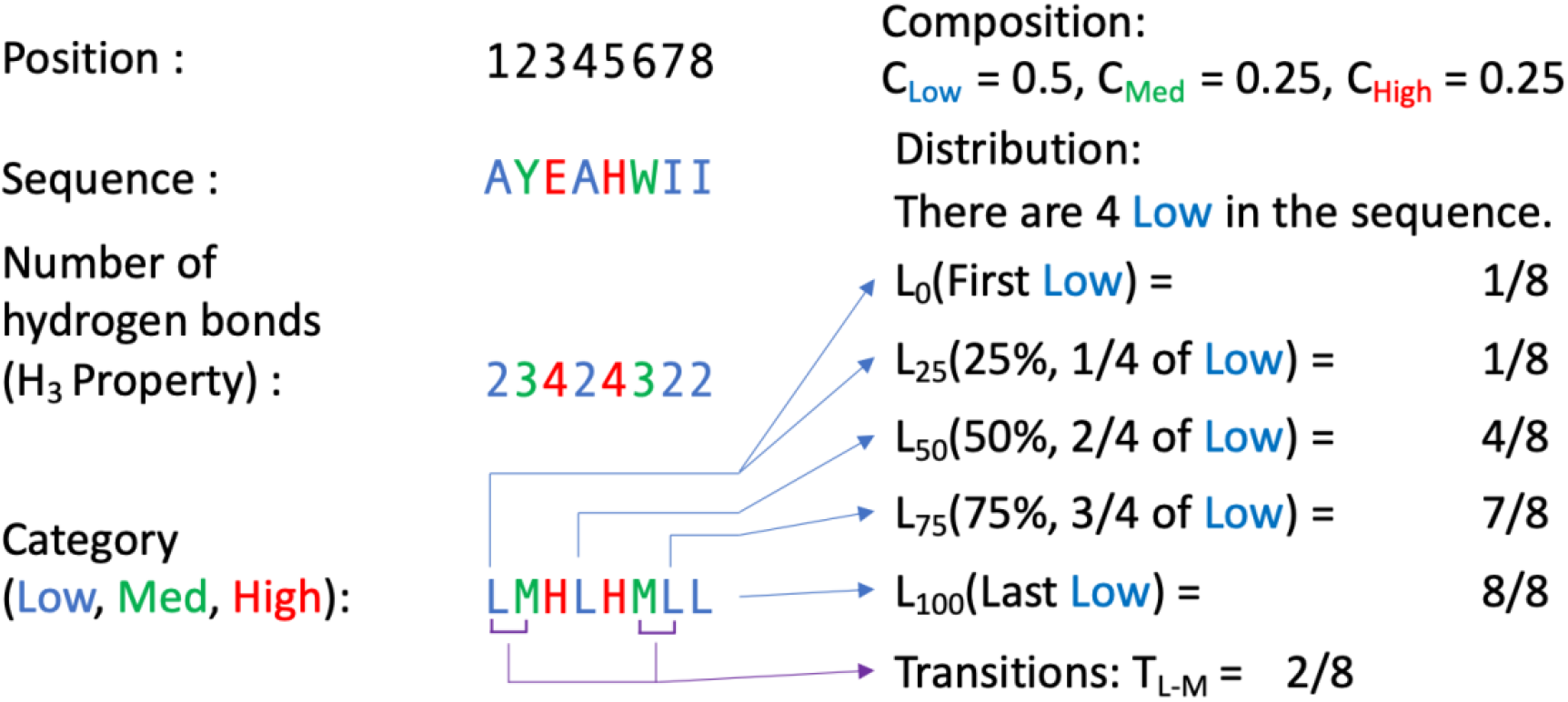
Deriving the physicochemical features: example of “number of hydrogen bonds”. In the toy sequence AYEAHWII, the “number of hydrogen bonds” feature can take values of 2, 3, or 4 at each amino acid. Translating each amino acid to its corresponding value and then to its category yields 23424322 and LMHLHMLL. Here we show how the property-based features *composition*, *transitions* (L-M case: Low to Medium and Medium to Low), and *distribution* (Low case) are calculated. These and the remaining physicochemical property features (*transitions* for M-H and L-H and *distributions* for Medium and High cases) are computed in the same manner.

**Figure 2:**
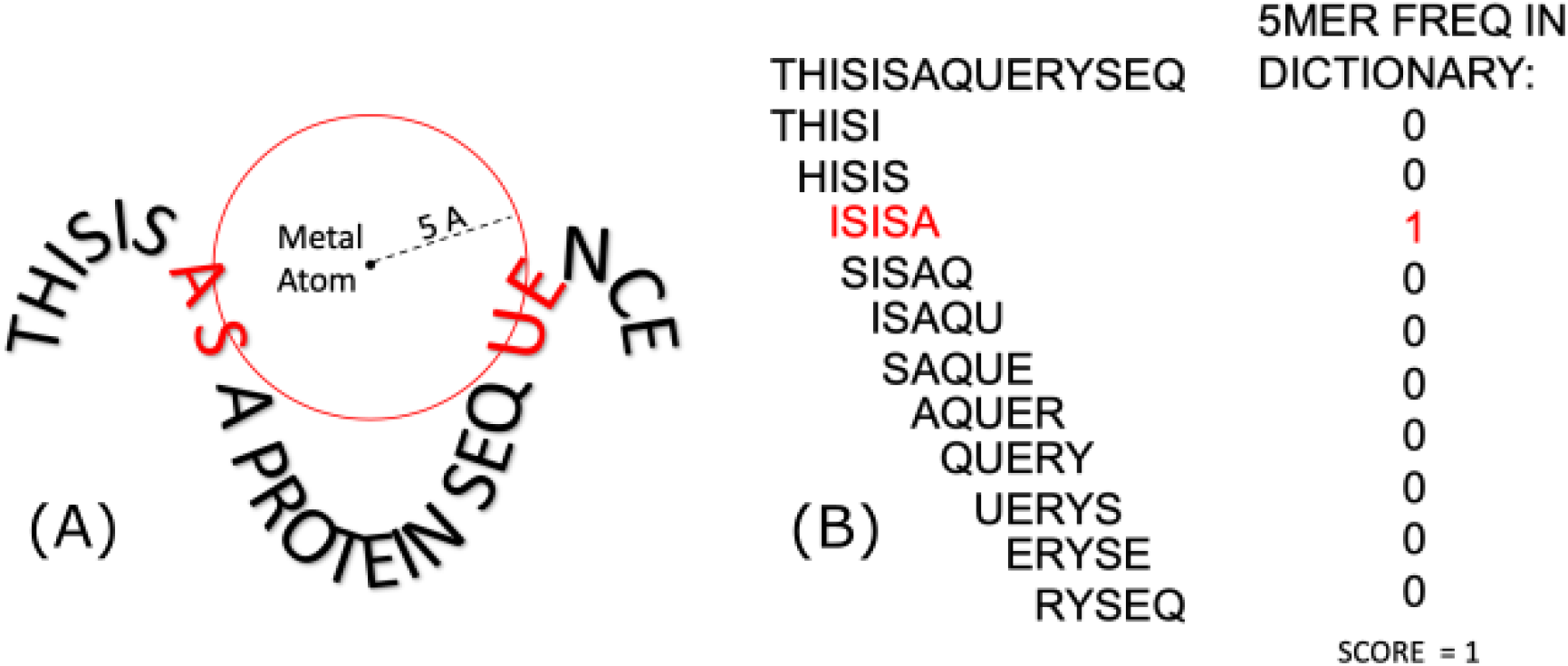
Deriving the 5mer features. In this toy example of a protein structure that binds a metal, the red circle depicts a sphere of 5Å radius around the metal; the residues within the red circle are marked red. Each of these red residues and their neighboring residues (two on each side) make up a feature 5mer. In this example, the four 5mers are: “ISISA”, “SISAP”, “EQUEN” and “QUENC”. The query sequence (right panel) is decomposed to count the number of metal-binding 5mers present. The final score for this feature is the sum of the counts of all 5mers in the sequence.

### Machine Learning

Using the above features, we trained a feed-forward Multi-Layer Perceptron (MLP) with back-propagation (BP) using the Keras [54] implementation in the machine learning framework Tensorflow [55]. Our model is a sequential network with the RMSprop [56] optimizer and a learning rate (lr)=0.000005. The optimization of the learning rate parameter was done via gradient descent, *i.e.* starting with an lr=0.5 we reduced it by an order of magnitude in each iteration of training and set the value to the one that minimizes the loss (calculated as binary cross-entropy). All other parameters were set at default values according to the Keras manual [57]. The input layer consists of 219 nodes – one node per feature. There are two hidden layers, as these are sufficient to approximate most partition problems and require considerably less computational power than more hidden layers [58]. Each layer had 219 nodes with a rectified linear unit activation function (or “Relu”) and a dropout of 0.2. Finally, there is single node output layer, using the sigmoid activation function and a default prediction (yes/no) cutoff set at 0.5.

We trained and tested our model for identifying metal-binding proteins using ten-fold cross-validation as follows: (1) we clustered sequences at 70% identity using CD-HIT and used the representative sequences of each cluster for training; (2) we split the resulting sequences into positives (metal binding) and negatives (nonbinding), and further divided each set into ten equally populated groups; (3) we built ten models by rotating through the ten splits using one positive and one negative group for testing and training with the other nine positive and negative groups. Since negatives in our set are more frequent than positives, we balanced the sets by randomly down-sampling the negatives to the same number as the positives for each model training. Note that the ten models were used to estimate the performance of the method, while the final *mebipred* model was constructed using all sequences and established parameters.

We additionally trained individual models with the same set of features to predict the binding of specific metals. We used the same modeling procedure and parameters as described above, only adding one more feature -- the score of the general metal-binding model above.

### Performance metrics

To measure the performance of our method we calculated overall accuracy, as well as positive precision, recall, and F-measure (all in Eqn. 1). True positives (TP) are metal-binding proteins predicted as metal-binding, false positives (FP) are metal non-binding proteins predicted as metal binding, false negatives (FN) are metal-binding proteins predicted as metal non-binding, and true negatives (TN) are metal non-binding proteins predicted metal non-binding.

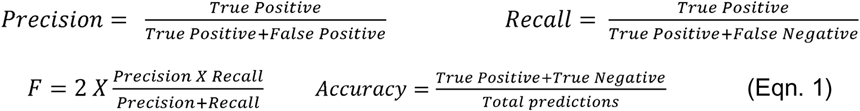

#### Comparing model performance to existing tools

To compare our method to a simple alignment-based approach, we extracted all sequences from the PDB. We generated a database of these sequences using the makeblastdb (−blastdb_version=5 and no extra parameters). We then ran BLAST (ncbi-blast+ V. 2.10.4) [60, 61] with default parameters (evalue 1; max_target_seqs 1000000) for all-to-all comparisons of protein sequences in this database. We used as gold standard our METAL and NO_METAL sets. For each e-value threshold, we counted the number of TP, FP, TN, and FN. Since we wanted to evaluate the use case where an unknown sequence is being annotated, we excluded self-hits from BLAST results but did not exclude hits to homologous sequences.

We further compared our performance to that of multiple published tools. For MetalDetector2 (Table 2), we used a set of non-redundant metal binding PDB structures described as the evaluation set of that method’s manuscript [50] (extracted in 2011). We also compared *mebipred* to two sequence based methods [40, 46] and a structure based method (MIB) [23] using the data from the BioLip database [62] (non-redundant at 90% sequence identity; Table 3).

### Generating short peptides

We took all 50-residue or longer protein sequences in the PDB (445,763 protein sequences) and, using a sliding window of 1, cut them into fragments of 50 residues (101,054,024 fragments).

### Metagenomic sample processing

To analyse metagenomic samples, reads were trimmed with trimmomatic [63] using default parameters. Trimmed reads were filtered with phred [64] using a score cut-off of 28. Reads were then analysed in two ways:

1. All reads where translated into the six possible reading frames using the standard bacterial codon table from Biopython [65]. Translated reads containing less than 15 amino acids where discarded. Remaining reads were used as input to *mebipred*
2. Reads were assembled using metaSPAdes [66] with variable kmer sizes (−k = 21,33,55,77,99, or 127). The resulting contigs were fed into Prokka [67] for ORF calling, gene annotation, and generation of protein sequence files. Prokka-annotated protein sequences were used as input to *mebipred*.

## Results and Discussion

### Available metal-binding protein structures are not diverse

The high resolution structure of most proteins is yet unknown, although this may change soon [68]. If a protein is of particular interest for the scientific community, it might be overrepresented in the PDB; e.g. over 1,300 structures of the SARS-COV2 spike protein. Thus, whether the known protein structures are a representative sample of all naturally occurring proteins is debatable and outside the scope of this work [69]. However, available structures constitute the most reliable set of metal-binding proteins [70, 71]. A quarter (49,996 of 152,346) of the PDB entries contain at least one of the metal atoms considered here. However, removing redundancy (at 70% sequence identity using CD-HIT [59]) retains 39,066 structures, of which only 9% are metal-binding (3,542 metal binding; note that a single sequence can bind multiple different ligands; SOM Data:PDB_chain_<METAL>_5.0A); the number of structures binding each ion varies (SOM Table 1),

**Table 1:**
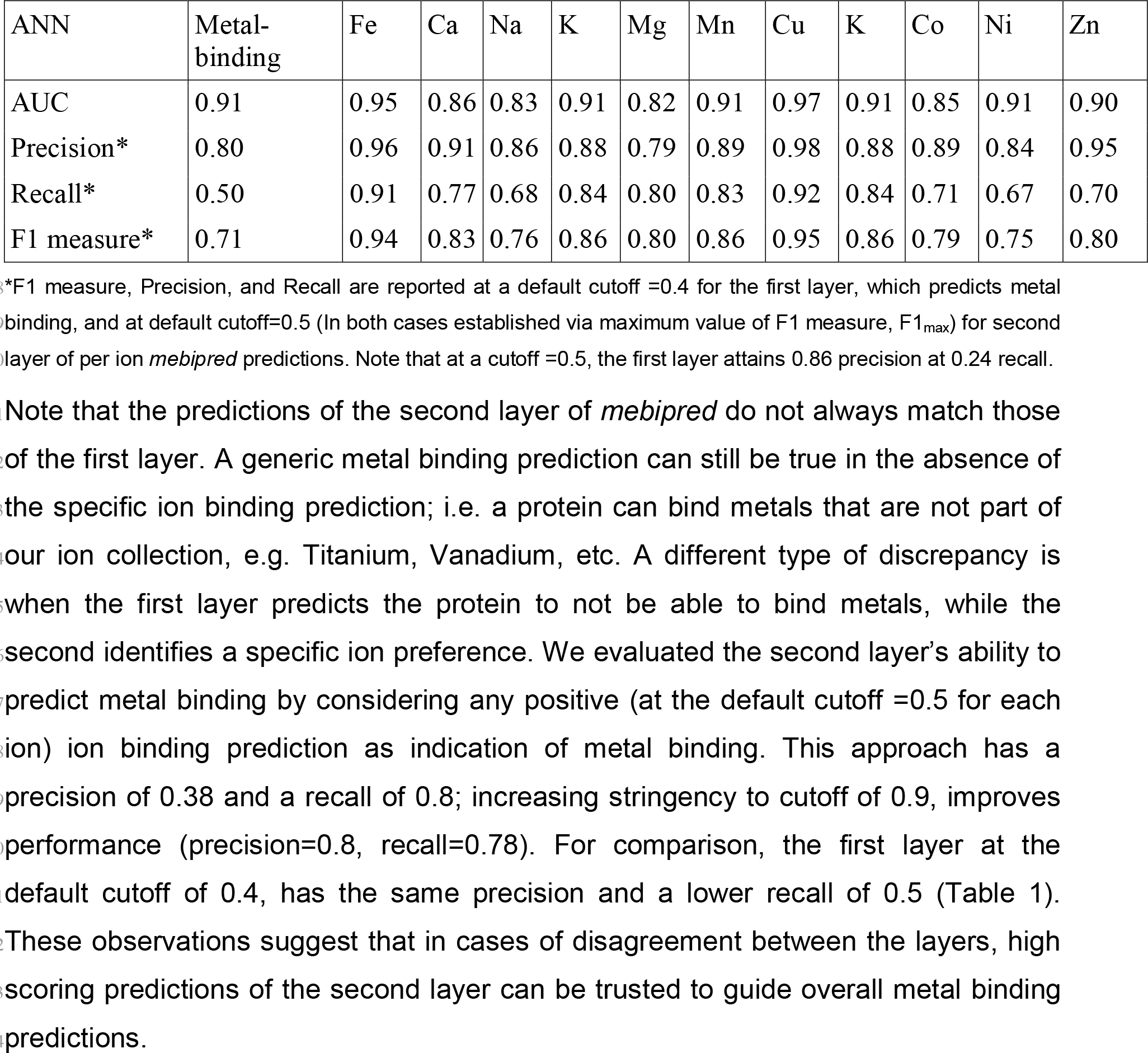
*mebipred* performance across metals.

| ANN | Metal-binding | Fe | Ca | Na | K | Mg | Mn | Cu | K | Co | Ni | Zn |
| --- | --- | --- | --- | --- | --- | --- | --- | --- | --- | --- | --- | --- |
| AUC | 0.91 | 0.95 | 0.86 | 0.83 | 0.91 | 0.82 | 0.91 | 0.97 | 0.91 | 0.85 | 0.91 | 0.90 |
| Precision* | 0.80 | 0.96 | 0.91 | 0.86 | 0.88 | 0.79 | 0.89 | 0.98 | 0.88 | 0.89 | 0.84 | 0.95 |
| Recall* | 0.50 | 0.91 | 0.77 | 0.68 | 0.84 | 0.80 | 0.83 | 0.92 | 0.84 | 0.71 | 0.67 | 0.70 |
| F1 measure* | 0.71 | 0.94 | 0.83 | 0.76 | 0.86 | 0.80 | 0.86 | 0.95 | 0.86 | 0.79 | 0.75 | 0.80 |

\*F1 measure, Precision, and Recall are reported at a default cutoff =0.4 for the first layer, which predicts metal binding, and at default cutoff=0.5 (In both cases established via maximum value of F1 measure, $F1_{max}$ ) for second layer of per ion *mebipred* predictions. Note that at a cutoff =0.5, the first layer attains 0.86 precision at 0.24 recall.

### *mebipred* attains exemplary performance

In cross-validation, the first (binary; yes/no) layer of *mebipred* identified sequence non-redundant metal-binding proteins with nearly 80% precision at a 50% recall (F1_max_=0.72 defines the default cutoff =0.4; Eqn. 1) – almost twice the precision obtained by BLAST at a similar recall (Fig. 3A); performance on the set of proteins including redundant sequences was, as expected, even higher (F1_max_ = 0.83 at default cutoff).

Note that performing the BLAST search for all sequences in the PDB took approximately six weeks (for 500,411 chain sequences in 152,346 structures), on average ~7.25 seconds/sequence on one core of a 2.4 GHz machine with 16G RAM). The same dataset was processed by *mebipred* on the same machine in 29 minutes (~6.3*10^−5^ seconds/sequence). While both BLAST and *mebipred* can be run with multiple cores, the difference in speed is likely to be retained. Moreover, BLAST compute time is expected to grow both with database size and the number of queries [72], while *mebipred* prediction time only reflects the number of queries, i.e. the algorithm scales as *(O)n*.

**Figure 3.**
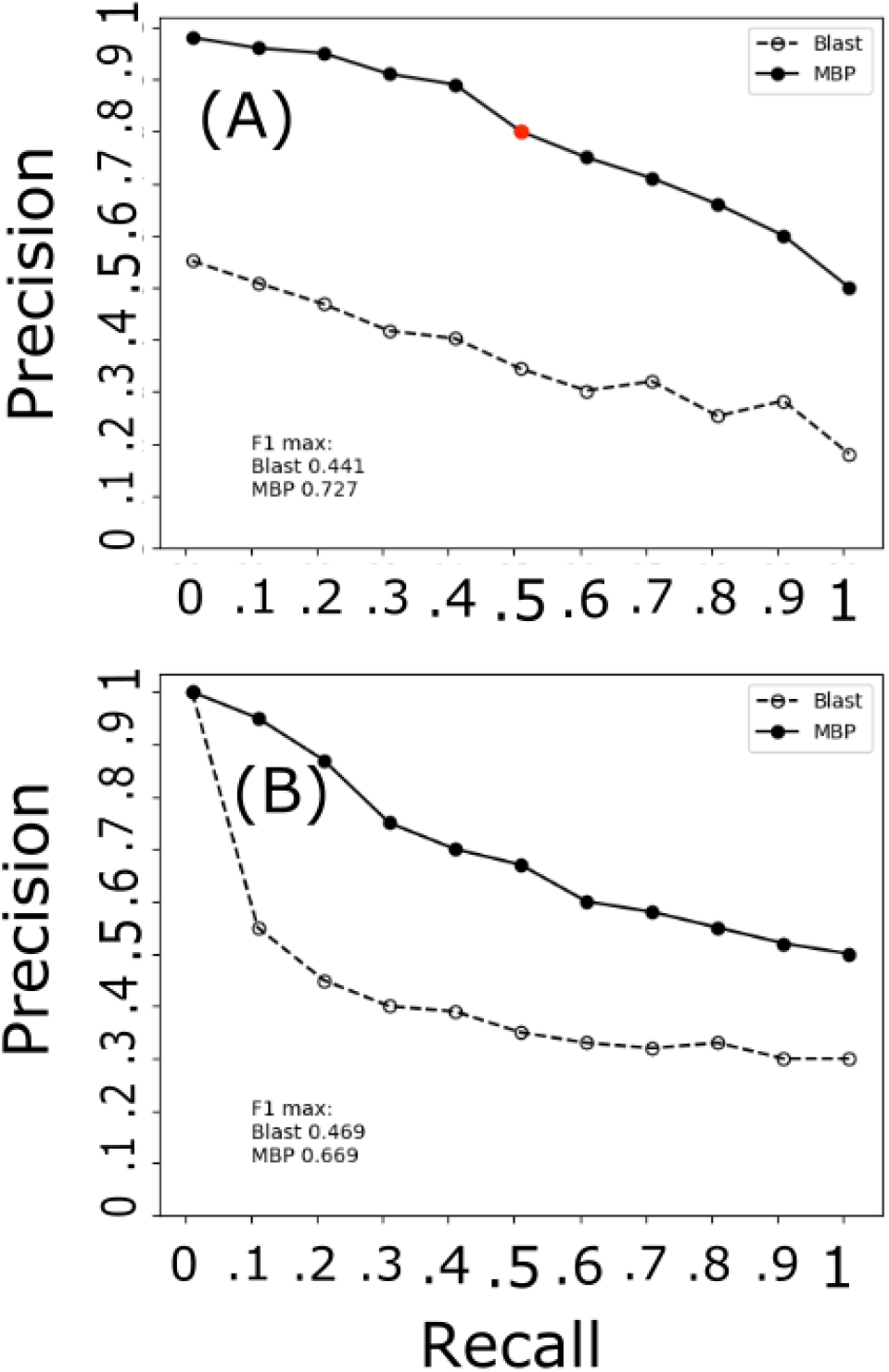
*mebipred* outperforms BLAST in identifying metal binding proteins and peptides. **(A)** At all cutoffs, *mebipred* (MBP; filled circles) is more precise than BLAST (empty circles). For example, at the default cutoff (score=0.4; red dot) it achieves 80% precision for half of the sequences (50% recall), as compared to 40% precision attained by BLAST. **(B)** *mebipred* also outperforms BLAST in identifying the metal binding propensity of proteins from their 50 amino acid fragment sequences. For example, for half of the fragments, it attains 67% accuracy, as compared to 39% attained by BLAST.

The second layer of *mebipred* predicts binding specifically to a ligand containing one of the ten ions under consideration. In cross-validation using our data set (Methods), *mebipred* was accurate in predicting ion specificity of individual proteins (Table 1). Note that we did not build predictors for proteins binding other biologically active metals (e.g. vanadium, molybdenum, titanium, etc.), because the number of structures binding these was insufficient to train a model of this kind. These could be incorporated into *mebipred* in the future if more protein structures binding these metals are resolved.

Note that the predictions of the second layer of *mebipred* do not always match those of the first layer. A generic metal binding prediction can still be true in the absence of the specific ion binding prediction; i.e. a protein can bind metals that are not part of our ion collection, e.g. Titanium, Vanadium, etc. A different type of discrepancy is when the first layer predicts the protein to not be able to bind metals, while the second identifies a specific ion preference. We evaluated the second layer’s ability to predict metal binding by considering any positive (at the default cutoff =0.5 for each ion) ion binding prediction as indication of metal binding. This approach has a precision of 0.38 and a recall of 0.8; increasing stringency to cutoff of 0.9, improves performance (precision=0.8, recall=0.78). For comparison, the first layer at the default cutoff of 0.4, has the same precision and a lower recall of 0.5 (Table 1). These observations suggest that in cases of disagreement between the layers, high scoring predictions of the second layer can be trusted to guide overall metal binding predictions.

Our evaluation of *mebipred* performance against that of other methods on our data was complicated by the absence of available web servers/standalone packages. Thus, we ran our tool on the data reported by the different methods. *mebipred* predicted metal-ligand binding better than Metal Detector2 [50] (Table 2) – a tool designed to predict transition metal-binding sites. Our method was also better than MetalExplorer [37], which predicts binding of eight metal ions with a precision of 60% at a range of recalls from 59% to 88%; *mebipred* attained an average of 80% precision for the same recall range. It also outperformed [40] (Table 3) and [46] (Table 3) methods, but performed worse than structure-based MIB [23]. Note that here we used the measure of accuracy (Eqn. 1) since it was reported in the corresponding publications, but precision and recall might be more relevant for imbalanced datasets [73].

**Table 2:**
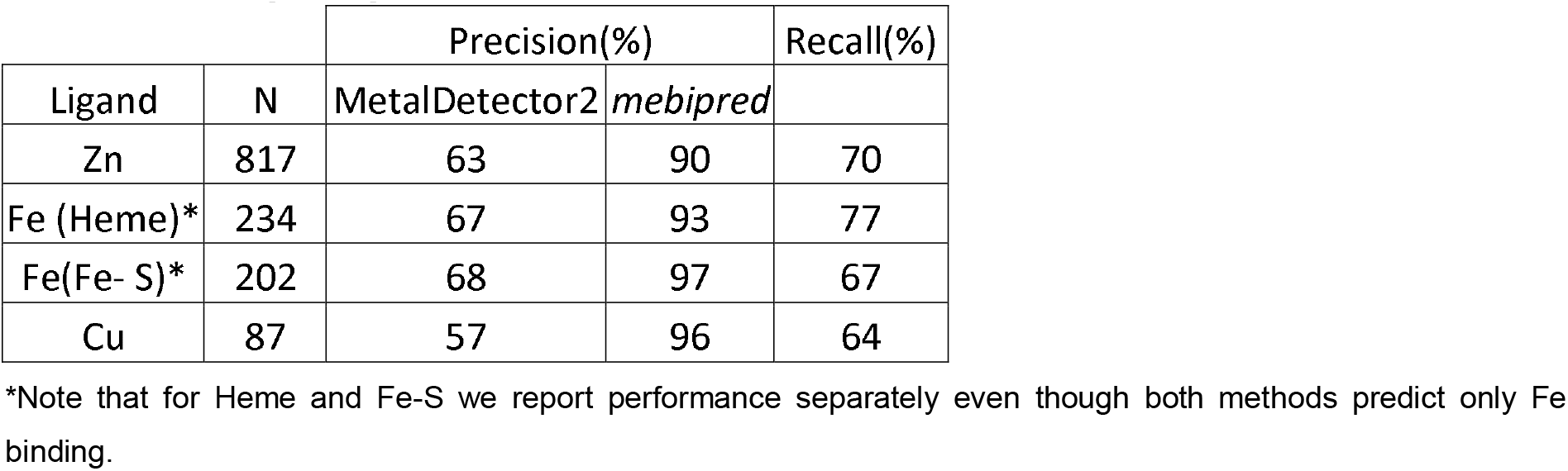
mebipred performance vs. MetalDetector2.

**Table 3:** mebipred performance vs. other methods.

| Ligand | Cao et al<br>[40]<br>(Acc%) | Kumar et al<br>[46] (Acc%) | MIB [23]<br>(Acc%) | *mebipred*<br>(Acc%) |
| --- | --- | --- | --- | --- |
|  | Sequence | Sequence | Structure | Sequence |
| Ca | 74.8 | 75.4 | 94.1 | 86.7 |
| Co | 83 | 85.3 | 94.7 | 86.2 |
| Cu | 96.3 | 78.1 | 95.3 | 87.2 |
| Fe2 | 91.3 | 75.6 | 95.1 | 89.2 |
| Fe3 | 87.8 | 74 | 94.9 | 89.2 |
| K | 80.3 | - | - | 74.0 |
| Mg | 75.3 | 74 | 94.6 | 75.6 |
| Mn | 83.2 | 68.8 | 95.0 | 89.7 |
| Na | 79.4 | 79.4 | - | 84.5 |
| Ni | - | 90.7 | 94.7 | 79.2 |
| Zn | 83 | 69 | 94.8 | 82.2 |

### *mebipred* predicts protein metal-binding propensity from short fragments

We extracted a set of 101,054,024 50-residue peptides from the PDB protein sequences (Methods); these correspond to the typical lengths of peptides that could be generated by translating DNA reads produced by next-gen sequencing [74]. We predicted metal-binding for these fragments using *mebipred* and aligned them (via BLAST) to PDB sequences following the same procedure as for complete proteins (Methods; excluding hits to proteins from which fragments were produced). *mebipred* outperformed BLAST (Fig 3B) in identifying peptides generated from metal-binding proteins. BLAST is not designed to deal with short sequence alignments [61, 75] and our results suggest that sequence identity may not be an accurate indicator of metal-binding either. Note that it is still possible that other alignment methods or changes in substitution matrices, *i.e.* penalizing the substitutions involving the residues often involved in metal binding, could yield better results.

### Ion binding preferences are consistent per Pfam family

We ran *mebipred* on the 607,903 Pfam proteins (8,207 families) whose structures are available in the PDB. According to binary predictions of the first layer, for 61% of the families, either all member proteins were predicted to be metal-binding or none were (SOM Data stats_with_id). Furthermore, for ~41% of the families, either all member proteins were predicted to bind a given metal or none were (SOM Data: stats_with_id). Of predictions per metal, 69% were cases where no members of one family bind that metal and 5% were cases where all of members of one family bind it – a total of 74% agreement of per ion predictions for members in the same family; the remaining 26% of the family members were predicted to have different ion preferences (SOM Data: stats_with_id). Our results indicate that metal binding preferences are mostly consistent within a Pfam family. This is expected, as Pfam domains reflect homology that, in turn, often suggests similar functionality and ligand binding [76, 77]. However, different ion preferences for a quarter of the families also suggest that specific metal availability within individual environments may have driven divergent evolution of new ligand-binding functionalities across organisms [22, 78]. Note that prediction error and cambialistic activity (i.e. ability to bind multiple ions) of certain proteins, which is not captured by this summary of ion binding, could also contribute to this discrepancy in metal binding preferences of single family members.

### *Mebipred* predictions do not always reflect existing annotations of metal-binding

We compared our METAL and NO_METAL datasets with Swiss-Prot metal-binding annotations. Of the 253,377 PDB sequences mapped to Swiss-Prot (PDBSWS [79]; April 2021), 53,652 (~20%) had annotations that disagreed with our data. Of these 32,667 were in our METAL set, *i.e.* in a PDB structure with a metal ion within 5Å of the chain (Methods) but were not described as metal binding by Swiss-Prot. Manual examination of ten randomly chosen discrepancies, confirms that the metal ion is present in a functional pocket, suggesting that Swiss-Prot annotations are incomplete. The remaining 20,985 sequences were described in Swiss-Prot as metal-binding but were not in our METAL set.

We ran *mebipred* on these 20,850 PDB–Swiss-Prot discrepancies. Our predictions (binary metal binding at default cutoff) agreed with Swiss-Prot annotations two thirds of the time (64%, 13,374 sequences, predicted metal-binding) sequences and with PDB otherwise (36%, 7,476 sequences, predicted metal non-binding). Crystal structures of metal-binding sequences may not contain a metal for a number of reasons, including biologically irrelevant binding (i.e. a metal can be bound by a protein, but isn’t under physiological conditions [80]) or experimental/technical crystallization decisions [16]. However, we expect that the 1,302 (6% of 20,850) non metal-binding chains from metal ion-containing PDB structures are most likely to be true non-binders of that ion. In fact, *mebipred* predictions for these proteins agreed with PDB 41% of the time (540 sequences predicted to be nonbinding) – a somewhat better agreement (vs 36%) than that for other designated metal non-binders.

A closer inspection further informs the reasons for database annotation differences. For example, 32 of the 540 predicted metal non-binding PDB chains map to the Rieske subunit of cytochrome BC1 – a Fe-S cluster binding protein (Swiss-Prot ID: Q5ZLR5) [81]. None of these 32 chains, however, are complete sequences of the protein and none contain the part of the structure that would bind the Fe-S cluster. In this particular case, the annotation discrepancy arises from a technical decision not to determine the metal-binding regions via crystallograhy [81]. While this level of scrutiny for every disagreement between databases is beyond the scope of this work, we note that an annotation discrepancy doesn’t necessarily constitute a “bug” but, rather, a feature of the method; i.e. *mebipred* could be used to resolve annotation conflicts between databases.

### *Mebipred* can predict metal-binding from metagenome read translations

We compared the metal-binding profiles of the Black Sea metagenomic (assembled and not assembled) samples obtained at different depths in a water column [82] (six samples (SRR12347146, SRR12347144, SRR12347141, SRR12347140, SRR12347142, SRR12347143, extracted from NCBI-SRA DB [83] and processed as in Methods). The relative frequencies of the resulting metal-binding protein/peptide predictions were very similar (Table 4; Euclidean distance between metagenome samples (*p* and *q*; Eqn. 2) =0, where *n* ∈ (Ca, Co, Cu, Fe, K, Mg, Mn, Na, Ni, Zn), indicates identical metal-binding frequency profiles).

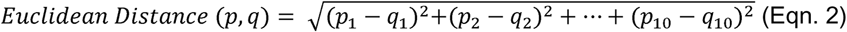

**Table 4:** Relative frequencies of metal binding proteins in metagenomic samples.

| SAMPLE (m) | Assembled |  |  |  |  |  |  |  |  | Euclidean distance |
| --- | --- | --- | --- | --- | --- | --- | --- | --- | --- | --- |
|  | Ca | Co | Cu | Fe | K | Mg | Mn | Na | Ni |  |
| 50 | 19.32 | 2.57 | 4.68 | 15.37 | 7.79 | 21.38 | 7.62 | 10.9 | 4.35 | 0.66 |
| 80 | 17.42 | 2.94 | 5.09 | 15.02 | 8.63 | 20.63 | 8.44 | 10.76 | 5.63 | 0.67 |
| 170 | 17.9 | 2.96 | 5.26 | 15.12 | 8.72 | 20.69 | 8.12 | 9.83 | 5.92 | 0.63 |
| 250 | 17.67 | 2.91 | 5.23 | 15.97 | 8.18 | 20.38 | 8.06 | 10.32 | 5.42 | 0.65 |
| 500 | 16.87 | 3.07 | 5.11 | 15.92 | 8.23 | 20.41 | 8.01 | 10.75 | 5.41 | 0.41 |
| 1000 | 16.91 | 2.95 | 5.11 | 16.4 | 8.28 | 20.37 | 7.96 | 10.51 | 5.4 | 0.47 |
| 2000 | 17.01 | 2.93 | 5.21 | 16.03 | 8.06 | 20.49 | 8.01 | 11.22 | 5.04 | 0.68 |

| *Unassembled* | % |  |  |  |  |  |  |  |  | Euclidean distance |
| --- | --- | --- | --- | --- | --- | --- | --- | --- | --- | --- |
| SAMPLE (m) | Ca | Co | Cu | Fe | K | Mg | Mn | Na | Ni |  |
| 50 | 19.04 | 2.60 | 4.70 | 15.60 | 7.59 | 20.89 | 7.65 | 10.77 | 4.24 | 0.66 |
| 80 | 17.50 | 2.92 | 5.04 | 14.93 | 8.78 | 21.17 | 8.29 | 10.44 | 5.68 | 0.67 |
| 170 | 18.24 | 2.94 | 5.32 | 15.41 | 8.83 | 20.97 | 7.98 | 10.12 | 5.91 | 0.63 |
| 250 | 17.63 | 2.83 | 5.24 | 15.89 | 8.07 | 19.82 | 8.24 | 10.10 | 5.37 | 0.65 |
| 500 | 16.81 | 3.08 | 5.15 | 15.86 | 8.01 | 20.16 | 8.15 | 10.67 | 5.25 | 0.41 |
| 1000 | 17.18 | 2.99 | 5.05 | 16.69 | 8.10 | 20.41 | 8.00 | 10.38 | 5.47 | 0.47 |
| 2000 | 17.23 | 2.97 | 5.11 | 15.70 | 7.91 | 21.01 | 7.95 | 11.16 | 5.01 | 0.68 |

This result suggests that *mebipred* can reliably predict metal-binding from translations of metagenomic reads (Methods).

### Diversity of metal-binding proteins highlights environmental differences

Across a number of environmental samples, we observed protein metal-binding signatures consistent with environmental features and subtypes.

#### Black Sea water column

From the above analysis we observed that the percentage of reads predicted as metal binding was approximately 1% for all Black Sea samples (SOM Table 4). The Black Sea is a heavily stratified body of water, where pH, oxygen, and light gradients have been characterized [84]. The sea surface layers where photosynthesis can occur, i.e. the epipelagic zone, are, by definition, up to 200m in depth; on the Black Sea, however, almost no photosynthetic activity can be found below 100m [85]. The epipelagic zone samples in our set are slightly enriched (2% increase) in Mg-binding proteins (Fig. 4). This is in line with the use of Mg in chlorophyll [86].

**Figure 4.**
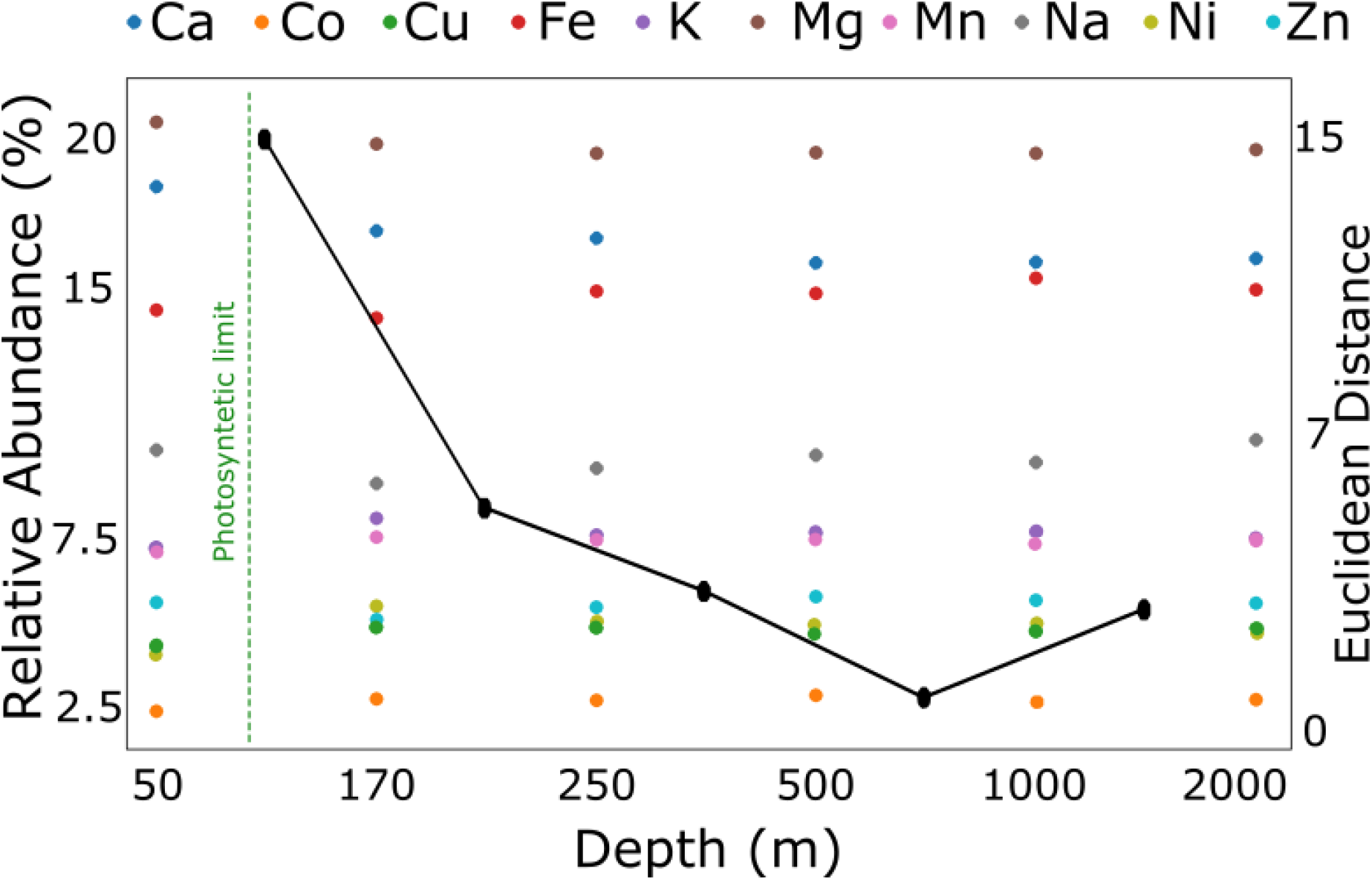
Predictions of metal binding proteins present in the Black Sea. The points on the graph indicate the relative abundance of ion-binding proteins (left y-axis) predicted from metagenomic samples collected at different depths of the Black Sea (x axis). The black line represents the Euclidean distance (right y-axis) between the vectors of predicted abundances at sequential depths. The markers on the line are placed between the depth measurements in each comparison. Samples show a phase transition (large Euclidean distance) at the photosynthetic limit (60 to 100m) {Gorlenko, 2005 #88;Callieri, 2019 #89}.

In non-photosynthetic environments, we observed a trade-off between the enrichment of Mg and Fe binding proteins, which can be accounted for by the lower pH increasing Fe availability and by the abundance of iron-reducing organisms at greater depths [87, 88]. The maximal difference between the abundances of predicted metal-binding proteins is observed between the samples taken at 50 and 170 meters, i.e. bypassing the photosynthetic limit; as indicated by the steep slope of the line tracing the Euclidian distance between metal-binding protein abundance vectors of individual samples (Fig. 4). Sample metal-binding preferences appear more similar below 170m (lower absolute value of slope). The difference between consecutive depths until 1,000m is in line with the changes in the environment described by the pH chemocline, changes in reduction potential, and reduced light [89]; i.e. the deeper one goes the lower the pH, the less calcium, and the more Fe [90]. The change in the sign of the slope indicating increasingly different samples at 1,000m and 2,000m likely accompanies a change in the microbial community [82]. This may reflect the transition from the Mesopelagic (200m to 1000m), where some light and oxygen are still available, to the Antropelagic region (1000m to 4000m), where there isn’t any of either. Alternatively, this change can highlight the fact that 2000m is essentially the seafloor [91].

#### Hot spring sediments

To test *mebipred* we analyzed 16 metagenomic samples from hot spring sediments obtained from NCBI-SRA DB (accession numbers SRS6512483) and previously described in Fullerton et al. [92]. The proportion of genetically encoded proteins binding each metal was similar (within 2%) for all samples (SOM Table 2). We observed a significant correlation between the relative frequency of proteins iron-binding iron and the iron environmental concentrations (Pearson r=0.54 p-val = 0.03; SOM Table 3); for zinc and manganese, the correlation was positive, but weak (Pearson r=0.1 and 0.18, p-val = 0.71 and 0.05, respectively). Copper and nickel binding proteins, on the other hand, had a negative correlation (but not significant) with the corresponding environmental concentrations (Pearson r=−0.1/p-val = 0.7 and Pearson r=−0.43/p-val=0.1, respectively) We lack complete information about metal requirements for different microbial strains. However, there is evidence that metabolites reflect the microbial community composition by altering the abundance of metabolite-relevant genes [93] – a finding somewhat in line with our observations. However, why did only iron (Fe) concentrations significantly correlate with iron-binding protein abundance? Fe is considered a major element (>1,000ppm), while others (Zn, Mn, Cu, Ni) are trace elements (<100 ppm) [94–96]. Metabolic requirements for each metal vary across organisms. However, iron is essential for nearly all of them; e.g., restricting iron availability to microbial invaders is part of the innate immune response [97]. Additionally, of the five measured metals, Fe is the only one that is present in the sampling sites at concentrations (observed: 3 – 400 ppm) below the what is needed for growth of metal requirement annotated bacteria [94, 96] (average requirement: 5,400 ppm); in fact, bacteria aim to actively accumulate Fe using specialized proteins [98]. The other four metals are usually required in concentrations [96] below those observed in this study. Moreover, higher concentrations may be deleterious to organism fitness, particularly for the anticorrelated metals. For example, nickel is required in trace quantities [99] and competes with Mg and Ca for binding sites [100]; in high concentrations, it can also damage DNA [101]. Copper is frequently toxic for bacteria at environmental concentrations [102] and is thus tightly regulated. Thus, given its key role in metabolism and limiting factor status, iron concentrations could drive microbial selection and explain the abundance of genes encoding iron binding proteins.

#### Human-host microbiomes

We further used *mebipred* to analyse randomly chosen human host and soil microbiome samples from the NCBI-SRA DB (SRR13422487, SRR12347145, ERR2855085, ERR5056238, SRR13422469, SRR12347144, SRR12432127, SRR12347141, SRR12360453, SRR12347140, SRR12347142, ERR2855082, SRR12347143, SRR12432116, SRR12347146, ERR2728283). Predicted metal-binding proteins (Fig. 5) are in line with the available metals in each environment. For example, few or no iron-binding proteins are predicted in samples of human origin except for one vaginal sample, where the occurrence may be explained by menstrual cycle bleeding. Low concentrations of iron-binding proteins are observed in the gut and pregnancy-associated vaginal microbiota, both of which may be accounted for by minor bleed episodes. As mentioned above, iron sequestering is part of normal human immune response and is lethal to most pathogenic bacteria [97]; normal non-pathogenic microbiota are likely to be adapted to low iron environment [103].

**Figure 5:**
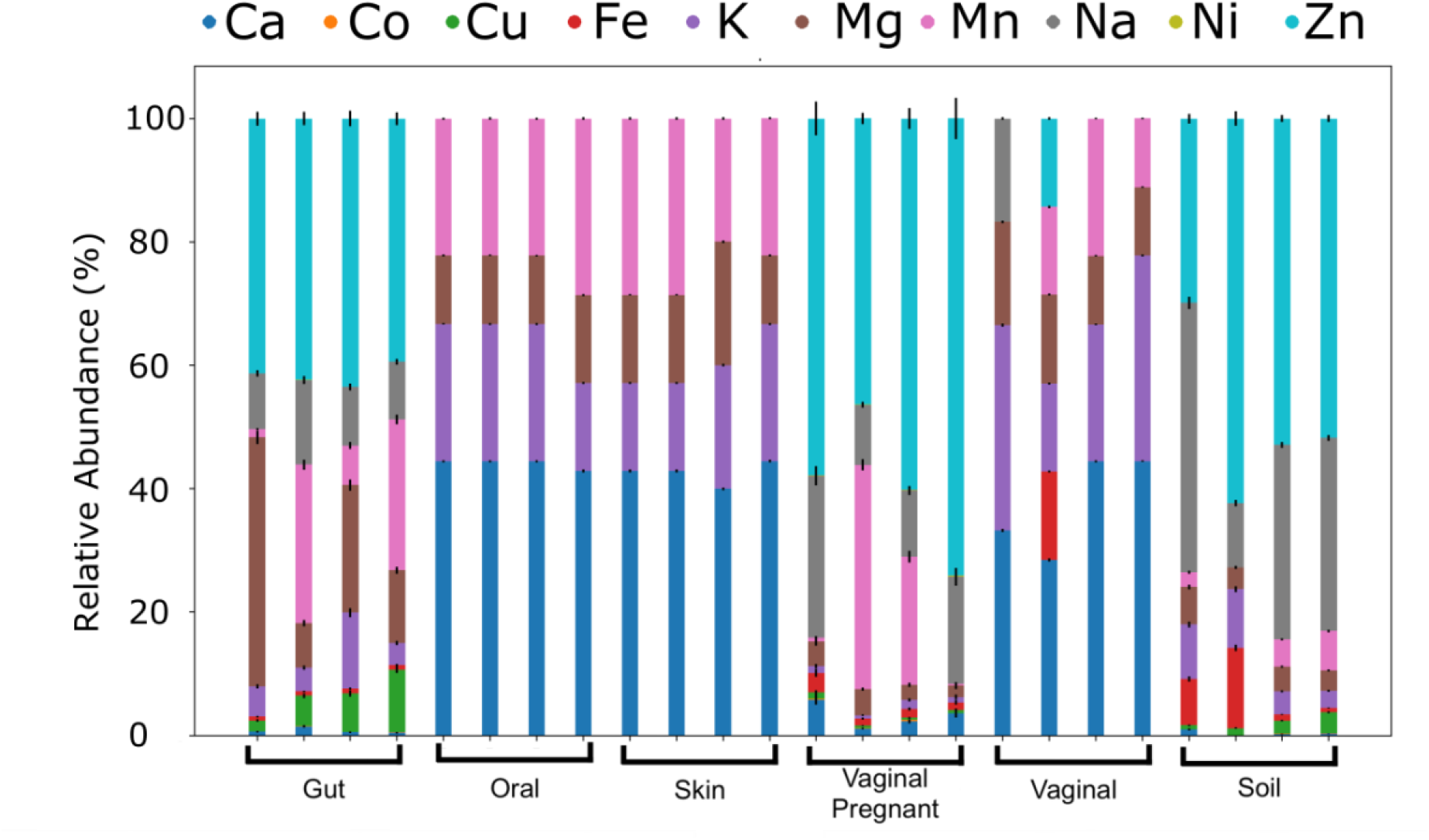
Differential abundance of metal binding proteins across environments. Each bar represents the relative abundance of predicted metal-binding proteins (y-axis) in a given metagenomic sample; four sampled per environment (x-axis). Concentration of these proteins per environment (column colors and sizes) are similar within and different across environments, suggesting signature metal ion preferences.

Metal-binding proteins predicted to occur in the soil and in gut samples target more different metals than do skin, mouth, and vaginal samples, likely due to the metabolic diversity of the former [104, 105]. The predicted metal-binding proteins in skin samples target metals (Ca, K, Mg, Mn) that can be found in sweat in relatively high concentrations (>1mg/l) [106]. Other metals (e.g. Zn, Cu) are present in sweat in trace concentrations (<1mg/l) [107, 108] and, consequently, few proteins bindings these metals are predicted (<1% of predictions). Furthermore, the differences in metal-binding protein abundances between vaginal samples from pregnant and non-pregnant women could reflect the large changes in the vaginal microbiome associated with pregnancy [109].

*mebipred* is an advance in the field of function prediction from protein sequence, which we showed to be applicable to the annotation of metagenomic samples. It can help resolve database annotation errors and shows potential for linking function with environmental conditions. We further expect that as more metal-binding protein structures are resolved, our method can be improved and expanded, for example to the detection of to other metals not currently treated. Its capacity to annotate metal binding informs the descriptions of microbiome environmental conditions and diversity. Finally, since most enzymes are metal binding proteins, it could also help enzyme prospecting in future studies.

## Conclusion

Here we compiled a gold-standard experimentally-derived metal-binding protein set and built *mebipred* – a sequence-based neural network predictor of metal binding. *mebipred* significantly outperforms existing sequence-based methods for annotation of metal binding; it also detects specific metals bound by each protein. We expect that the growth in the number of metal binding proteins with resolved structures will make these types of approaches even more powerful in the near future. To the best of our knowledge, *mebipred* is also the only reference-free sequence-based tool for identifying metal-binding. Our method is faster than existing tools and can predict metal binding using short protein fragments – both characteristics that make it useful in analysis of metagenomic data. In evaluation of microbiome samples we found that differences in the number of predicted metal-binding proteins were related to the concentration of metal ions in the corresponding environments.

## Abbreviations used

*mebipred*: Metal-binding predictor
PDB: Protein Data Bank
BLAST: Basic local alignment search tool

## Acknowledgments

We are grateful to Drs. Paul Falkowski, Maximilian Miller, Chengsheng Zhu, Yannick Mahlich, Kenneth McGuinnes, Adrienne Hoarfrost, Natalia Rigazio, and Zishuo Zeng (all Rutgers) for their critical comments on the development of the method. We would also like to express gratitude to the PDB team and to all researchers that solve and deposit protein structures into the PDB for nearly half a century. Without them this work, and most of other structural bioinformatics work, would not be possible.

DG has received funding from the European Research Council (ERC) under the European Union’s Horizon 2020 research and innovation programme (Grant agreement No. 948972) ERC-STG-2020 project CoEvolve. DUF is a fellow of the CONICET. Y.B. and A.A. were supported by the NASA Astrobiology Institute grant 80NSSC18M0093. Y.B. was also supported by the NSF (National Science Foundation) CAREER award 1553289.

## Notes

### Competing Interest Statement

The authors have declared no competing interest.

