## Supplementary material for "mebipred: identifying metal-binding potential in protein sequence": Suplemental tables

Supporting online material
for:
mebipred: identifying metal-binding abilities from protein sequence.

Aptekmann AA^1,2*^, Buongiorno J^5^, Giovanelli D^124^, Glamoclija M^6^ , Ferreiro DU^3^, Bromberg Y^1,*^

Table of Contents for Supporting Online Material

Short description of Supporting Online Material

Material

| ion | 5A_PDB | 5A_PDB70 |
| --- | --- | --- |
| Ca | 10486 | 1606 |
| Co | 1130 | 131 |
| Cu | 1447 | 167 |
| Fe | 7151 | 964 |
| K | 1040 | 59 |
| Mg | 10474 | 1415 |
| Mn | 2408 | 248 |
| Na | 5275 | 790 |
| Ni | 1272 | 200 |
| Zn | 9672 | 1707 |
| Total | 50355 | 7287 |

Supplementary Table 1: number of structures binding each ion in the PDB and PDB70 (redundancy reduced at 70% sequence identity) databases.

| Sample | Ca | Co | Cu | Fe | K | Mg | Mn | Na | Ni | Zn |
| --- | --- | --- | --- | --- | --- | --- | --- | --- | --- | --- |
| 1 | 15266 | 2406 | 4656 | 11062 | 10947 | 18544 | 8066 | 9368 | 8459 | 4479 |
| 2 | 16315 | 2653 | 4831 | 12687 | 12400 | 20825 | 9217 | 10817 | 10124 | 5555 |
| 3 | 70528 | 11442 | 21944 | 52321 | 54663 | 87904 | 39892 | 48720 | 43242 | 20696 |
| 4 | 17217 | 2810 | 4926 | 13957 | 12540 | 22162 | 9056 | 10136 | 10171 | 6471 |
| 5 | 27799 | 4642 | 8670 | 20903 | 20750 | 34372 | 15629 | 18244 | 16430 | 8401 |
| 6 | 43900 | 6967 | 13124 | 32821 | 31516 | 52025 | 23236 | 27547 | 24516 | 12728 |
| 7 | 9004 | 1522 | 2728 | 6959 | 6682 | 11635 | 4958 | 5696 | 5220 | 2795 |
| 8 | 21620 | 3207 | 6050 | 14930 | 15430 | 25819 | 11488 | 13037 | 11614 | 6245 |
| 9 | 30904 | 4654 | 9007 | 22113 | 22431 | 37631 | 16327 | 18946 | 17347 | 9431 |
| 10 | 16244 | 2338 | 4585 | 11993 | 12105 | 20108 | 8788 | 10166 | 9463 | 5221 |
| 11 | 41958 | 6551 | 12330 | 30652 | 31579 | 52692 | 23292 | 27326 | 24914 | 13189 |
| 12 | 30830 | 5653 | 9601 | 24851 | 24426 | 41000 | 18344 | 21501 | 20387 | 10008 |
| 13 | 66204 | 10806 | 20108 | 51642 | 50031 | 83913 | 36891 | 43449 | 41416 | 21723 |
| 14 | 9020 | 1557 | 2978 | 7225 | 7125 | 11678 | 5463 | 6355 | 5901 | 2976 |
| 15 | 16376 | 2693 | 4917 | 11999 | 12811 | 20867 | 9406 | 11525 | 10197 | 4978 |
| 16 | 10862 | 1933 | 3349 | 8782 | 8451 | 14443 | 6037 | 7167 | 6737 | 3498 |

Supplementary Table 2: Predicted metal binding protein for each marine sample.

| Sample | Cu | Fe | Mn | Ni | Zn |
| --- | --- | --- | --- | --- | --- |
| 1 | 3.12 | 3.97 | 11.43 | 0.43 | 2.28 |
| 2 | 0.48 | 6.55 | 39.34 | 0.16 | 13.85 |
| 3 | 1.24 | 0.63 | 9.80 | 0.16 | 0.70 |
| 4 | 1.15 | 2.80 | 12.90 | 0.62 | 2.54 |
| 5 | 3.62 | 1.05 | 58.20 | 0.62 | 1.55 |
| 6 | 1.30 | 4.72 | 21.58 | 0.35 | 1.35 |
| 7 | 0.94 | 0.06 | 0.14 | 0.02 | 0.10 |
| 8 | 1.63 | 1.24 | 13.34 | 1.26 | 1.63 |
| 9 | 2.20 | 1.87 | 31.62 | 1.91 | 2.24 |
| 10 | 2.54 | 2.22 | 28.22 | 2.04 | 2.22 |
| 11 | 1.10 | 1.17 | 18.06 | 0.96 | 1.49 |
| 12 | 0.17 | 7.34 | 55.70 | 0.03 | 0.24 |
| 13 | 0.17 | 7.34 | 55.70 | 0.03 | 0.24 |
| 14 | 0.89 | 1.20 | 19.79 | 0.30 | 1.51 |
| 15 | 1.63 | 1.24 | 13.34 | 1.26 | 1.63 |
| 16 | 4.06 | 2.15 | 38.02 | 0.59 | 8.19 |

Supplementary Table 3: Ion concentrations for each sample (mM)

| Sample | Ca | Co | Cu | Fe | K | Depth(m) | Reads | mbp/n_reads% |
| --- | --- | --- | --- | --- | --- | --- | --- | --- |
| SRR12347146 | 1105 | 146 | 264 | 872 | 446 | 50 | 1018328 | 0.56% |
| SRR12347144 | 7652 | 1270 | 2249 | 6484 | 3763 | 170 | 4382497 | 0.98% |
| SRR12347143 | 8009 | 1327 | 2363 | 7230 | 3706 | 250 | 4392416 | 1.03% |
| SRR12347142 | 6738 | 1217 | 2035 | 6355 | 3259 | 500 | 4103257 | 0.97% |
| SRR12347141 | 10831 | 1873 | 3251 | 10453 | 5284 | 1000 | 5967681 | 1.07% |
| SRR12347140 | 9429 | 1631 | 2920 | 8955 | 4507 | 2000 | 4153399 | 1.34% |
| Sample | Mg | Mn | Na | Ni | Zn | Depth(m) | Reads | mbp/n_reads% |
| SRR12347146 | 1223 | 436 | 621 | 248 | 348 | 50 | 1018328 | 0.56% |
| SRR12347144 | 8864 | 3472 | 4227 | 2517 | 2339 | 170 | 4382497 | 0.98% |
| SRR12347143 | 9234 | 3656 | 4684 | 2497 | 2657 | 250 | 4392416 | 1.03% |
| SRR12347142 | 8078 | 3167 | 4279 | 2153 | 2478 | 500 | 4103257 | 0.97% |
| SRR12347141 | 13033 | 5099 | 6701 | 3448 | 3920 | 1000 | 5967681 | 1.07% |
| SRR12347140 | 11464 | 4481 | 6289 | 2812 | 3342 | 2000 | 4153399 | 1.34% |

Supplementary Table 4: Metal binding predictions for Black Sea samples.

| **Sample** | **Code** | **SRA** |
| --- | --- | --- |
| 1 | PFS | SRS6512461 |
| 2 | FAS | SRS6512462 |
| 3 | PGS | SRS6512467 |
| 4 | SLS | SRS6512479 |
| 5 | ETS | SRS6512459 |
| 6 | QNS | SRS6512473 |
| 7 | PLS | SRS6512468 |
| 8 | QHS2 | SRS6512470 |
| 9 | RSS | SRS6512475 |
| 10 | EPS | SRS6512493 |
| 11 | SIS | SRS6512481 |
| 12 | BRS1 | SRS6512487 |
| 13 | BRS2 | SRS6512485 |
| 14 | CYS | SRS6512489 |
| 15 | QHS1 | SRS6512471 |
| 16 | TCS | SRS6512483 |

Supplementary Table 5: SRA accessions corresponding to each sample.

| **Sample** | **Cu** | **Fe** | **Mn** | **Ni** | **Zn** |
| --- | --- | --- | --- | --- | --- |
| BRS1 | 10.99 | 403.71 | 3286.53 | 1.61 | 15.73 |
| BRS2 | 10.99 | 403.71 | 3286.53 | 1.61 | 15.73 |
| CYS | 55.87 | 66.22 | 1167.52 | 17.25 | 97.92 |
| EPS | 159.93 | 122.19 | 1665.15 | 118.06 | 144.40 |
| ETS | 227.86 | 57.59 | 3434.01 | 36.01 | 100.59 |
| FAS | 30.21 | 360.48 | 2320.79 | 9.23 | 900.45 |
| PFS | 196.57 | 218.30 | 674.09 | 25.14 | 148.44 |
| PGS | 78.41 | 34.40 | 578.45 | 9.41 | 45.49 |
| PLS | 59.00 | 3.30 | 8.01 | 1.17 | 6.60 |
| QHS1 | 102.48 | 68.03 | 787.16 | 72.87 | 106.12 |
| QHS2 | 102.48 | 68.03 | 787.16 | 72.87 | 106.12 |
| QNS | 81.80 | 259.85 | 1273.41 | 20.20 | 87.70 |
| RSS | 138.49 | 102.91 | 1865.37 | 110.80 | 145.92 |
| SIS | 69.53 | 64.26 | 1065.70 | 55.82 | 96.57 |
| SLS | 72.19 | 153.78 | 761.27 | 36.04 | 165.33 |
| TCS | 255.49 | 118.14 | 2243.36 | 34.34 | 532.13 |
| ARS | 142.25 | 55.19 | 1127.73 | 40.71 | 80.82 |

Supplementary Table 6: Ion concentrations for each sample (ppm)

References for Supporting Online Material
